# A comparison of ancestral state reconstruction methods for quantitative characters

**DOI:** 10.1101/037812

**Authors:** Manuela Royer-Carenzi, Gilles Didier

**Affiliations:** Aix-Marseille Université, CNRS, Centrale Marseille, I2M, UMR 7373, 13453 Marseille, FRANCE

**Keywords:** Ancestral state reconstruction, Maximum likelihood, Brownian motion, Energy distance

## Abstract

Choosing an ancestral state reconstruction method among the alternatives available for quantitative characters may be puzzling. We present here a comparison of five of them, namely the maximum likelihood, restricted maximum likelihood, generalized least squares, phylogenetic independent contrasts and squared parsimony methods.

A review of the relations between these methods shows that the first three ones infer the same ancestral states and can only be distinguished by the distributions accounting for the reconstruction uncertainty which they provide.

The respective accuracy of the methods is assessed over character evolution simulated under a Brownian motion with (and without) drift. We start by giving the general form of ancestral state distributions conditioned on leaf states under the simulation model.

Ancestral distributions are used first, to give a theoretical lower bound of the expected reconstruction error, and second, to develop an original evaluation scheme which is more efficient than comparing the reconstructed and the simulated states.

Our simulations show that: (i) the methods do not perform well as the evolution drift increases; (ii) the maximum likelihood method is generally the most accurate and (iii) not all the distributions of the reconstruction uncertainty provided by the methods are equally relevant.

## Introduction

Besides being essential to understand the process of character evolution, ancestral state reconstruction plays an important role in the study of ecological diversification and comparative analysis. Though it may concern more or less complex traits (either ecological, phenotypic, or biogeographic), we focus here on quantitative characters, i.e. measured as continuous variables such as weight, size etc.

From a methodological point of view, ancestral state reconstruction is a challenging problem which has been addressed by several approaches. The general question can be stated as follows. Taking as inputs the phylogeny of a set of organisms (given as a tree with branch lengths) and their character states, a reconstruction method has to infer - as accurately as possible - the character states of the ancestral organisms. The reconstruction approaches fall into two major classes: methods based on the parsimony principle (Fitch 1971; Swofford and Maddison 1987; Maddison 1991; Collins et al. 1994), whose general idea is to impute the missing values of the tree by minimizing the sum of distances between ancestors and their direct descendant characters, and methods based on stochastic models of character evolution, mainly Brownian motion for continuous traits (Schluter et al. 1997; Pagel 1999; Huelsenbeck and Ronquist 2001; Nielsen 2002). Several authors discuss the advantages of stochastic approaches over parsimonious ones (Schluter et al. 1997; Mooers and Schluter 1999; Pagel 1999; Nielsen 2002; Huelsenbeck et al. 2003). An important point is that stochastic approaches take into account divergence times (branch lengths) while parsimonious methods do not. Moreover, stochastic approaches may provide probability distributions of the reconstructed ancestral states, accounting for their uncertainty and which can be used to develop hypothesis testing and confidence intervals.

In our study, we focus on five widely-used reconstruction methods, namely the maximum likelihood, restricted maximum likelihood, generalized least squares, phylogenetic independent contrasts and squared parsimony methods. Before comparing their accuracy, we review the methods and their relationship to each other. It turns out that the first three ones reconstruct the same ancestral states. These three methods may still be distinguished, and to some extent compared, since they provide different probability distributions of their uncertainty.

Evaluating the respective performances of these methods is a natural and important question. Works aiming at answering this question proceed by comparing the reconstructed states with reference “trusted” ones. Such reference values for ancestral states may be obtained either by considering fossil character states or by simulating, via a stochastic model, artificial evolution of the character and by keeping track of the ancestral states observed during simulations (Martins 1999). Webster and Purvis (2002) and Oakley and Cunningham (2000) assess several reconstruction methods with regard to measurements on fossils. They both observe that the methods are confounded by an evolutionary trend toward increasing size.

Our comparison of the five methods is based on artificial evolution simulated under Brownian motions with and without drift. The artificial evolution runs on the phylogenetic tree of Pleistocene planktic Foraminifera (Webster and Purvis 2002). Besides the fact that we consider evolution models with drift, a noticeable difference with previous works is that the reconstructed states are compared with regard to the ancestral state distributions conditioned on the simulated leaves, rather than with the simulated ancestral states as it is done usually. Intuitively, in this way, we compare the reconstructed state with all the possible realizations of the evolution process with the given simulated leaf states. Moreover the ancestral distribution conditioned on the leaves does reflect the uncertainty inherent to the stochastic character of evolution as modeled in simulations. In particular, it allows us to determine a lower bound of the expected reconstruction error as well as the reconstructed state achieving this lower bound. This can be seen as a transposition of ideas of (Steel and Sz´ekely 1999) and (Royer-Carenzi et al. 2013).

Another motivation of this work is to assess the relevance of the distributions provided by the methods for the reconstruction uncertainty. These distributions are expected to provide a greater amount of information than single values for ancestral states (Schluter et al. 1997; Polly 2001). Altogether with our new comparison scheme, we compare the conditional ancestral distributions given the leaves with the distributions provided by the methods. A distance between distributions, called the Energy distance offers us a consistent framework to compare both reconstructed states and reconstructed probability distributions, with ancestral state distributions conditioned on leaves (Sz´ekely and Rizzo 2013). The Energy distance is strongly related to the absolute bias.

Finally, we provide exact, matrix-based, implementations of some of the methods which were formerly based on numerical optimization algorithms. Our R-scripts have been incorporated into the reconstruct function of the ape R-package since version 3.2 (Paradis et al. 2004, https://cran.r-project.org/web/packages/ape/index.html).

The rest of the paper is organized as follows. In Section 1, we present two standard models of quantitative character evolution. Section 2 briefly describes the reconstruction methods and shows how they are related. Section 3 is devoted to our assessment protocol. We provide the form of the ancestral distributions conditioned on the leaf states under the simulation. These ancestral distributions are next used to define our evaluation protocol and to give a lower bound of the expected reconstruction error. In its final version, the protocol is based on the Energy distance between probability distributions, both for assessing the reconstructed states and the distribution provided by the methods. The results of our simulations are finally presented and discussed in Section 4.

## 1 Models of evolution for quantitative characters

### 1.1 Phylogenetic trees - Notations

In the standard ancestral character reconstruction problem, one assumes that the evolutionary history of the species is known and given as a rooted phylogenetic tree with branch lengths. Our typical tree contains *n* + 1 nodes (including leaves), among which *r* are internal nodes (excluding the root). By convention, the nodes are indexed in the following way:

- index 0 for the root,
- indices 1 to *r* for the other internal nodes,
- indices *r* + 1 to *n* for the leaves.

The nodes are numbered in such a way that if a node *j* descends from a node *i* then *j* > *i*. For any non-root node *j*, we put *p*(*j*) for the index of its direct ancestor and *τ_j_* for the length of the branch relying *p*(*j*) to *j*.

Let *X* be a random variable. We put *f_X_* for its density function and 𝔼(*X*) for its expectation.

### 1.2 Models of evolution

We make the standard assumptions that character evolves independently along the branches of the phylogenetic tree and that its evolution is homogeneous both through time and lineages. In order to actually model the evolution of a character, we first need to define an initial probability density *f*_*Z*0_ for the state of the root. In the simulation models, *f*_*Z*0_ is the degenerate density at a given value *z*_0_, i.e. our simulations all start from a given root state *z*_0_ which is a parameter of the model. The reconstruction methods assume an improper flat density as initial probability density *f*_*Z*0_ (i.e. *f*_*Z*0_(*x*) = 1 for all *x*).

#### 1.2.1 Brownian motion model

The *Brownian motion* (BM) model is the simplest stochastic process able to model the evolution of a quantitative character (Felsenstein 1985; Schluter et al. 1997). Under this model, evolution is neutral and governed by a rate parameter *σ* which accounts for its diffusion. Formally, along a branch of the tree, the stochastic process (*X_t_*) accounting for the character state has the form:

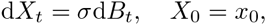

where (*B_t_*) denotes the standard Brownian motion, defined as a centered Gaussian process with stationary and independent increments, and *B_t_* ∽ *𝒩*(0,*t*), where *𝒩*(µ, σ^2^) is the Gaussian distribution of mean µ and variance *σ^2^*. Thus increments (*X_t+S_ 𝒩 X_t_*) are independent with law *𝒩* (0, σ^2^*s*).

#### 1.2.2 Arithmetic Brownian motion model

Biological evolution is not always assumed to be neutral. For instance, Cope’s rule states that species tend to increase in body size over time (Kingsolver and Pfennig 2004; Van Valkenburgh et al. 2004; Hone and Benton 2005). In Webster and Purvis (2002), fossil evidence suggests that a neutral process cannot model the evolution of Pleistocene planktic Foraminifera size since it tends to increase with time. A similar observation is made by Oakley and Cunningham (2000).

The *Arithmetic Brownian motion* (ABM), sometimes called Brownian motion with drift, yields to model a linear drift µ, which can be either positive or negative. Along a branch of the tree, the stochastic process (*X_t_*) of the character state now has the form:

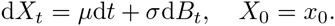

The increments (*X*_*t+s*_ — *X_t_*) are independent with law *𝒩* (*µs*, *σ*^2^ *s*).

Note that these two models are basically embedded: a **BM** is nothing but an **ABM** with drift-parameter *µ* = 0.

### 1.3 Likelihood of a character evolution

Let us consider a particular realization of the evolution process, which is known only through the vector (*z*_0_, *z*_1_, …, *z_n_*) of the character states at the nodes of the tree (i.e. entry *z_i_* is the character state at node *z_i_*). The increments of the character state between nodes and their children give us a natural expression of the likelihood of such a vector. Let us put *φ*(., *θ*^2^) for the density of a centered normal law with variance *θ*^2^:

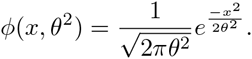

Under the **BM** model with parameter *σ^2^* and root probability density *f_Z_0__*, the likelihood of a realization (*z_0_, z_1_, …, z_n_*) is

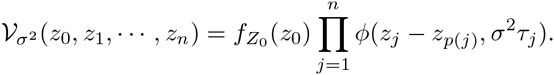

The corresponding log-likelihood is

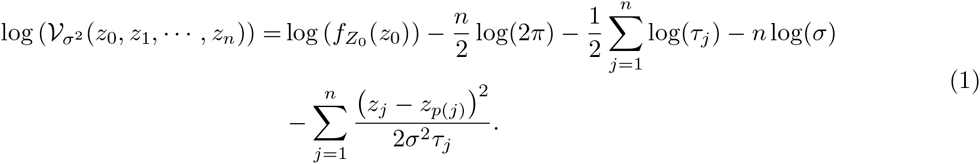

## 2 Reconstruction methods

### 2.1 Presentation

#### 2.1.1 Brownian-based methods

We present here four methods all relying on the assumption that the character evolves following a **BM** model with an improper flat distribution for the root state. Their current implementations return not only a reconstructed state for each internal node *j* but also a probability distribution of this quantity. Therefore, the reconstructed state may be seen as a random variable 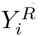, which accounts for the reconstruction uncertainty. Hereafter, we give a brief description of these four methods:

- The Maximum Likelihood method (*ML*) infers the ancestral states which maximize their joint likelihood under a **BM** model with an improper flat distribution for the root state (Schluter et al. 1997). This maximum likelihood estimation is simultaneously performed on the ancestral states and on the variance of the **BM** model. For any internal node *j, ML* returns the reconstructed state 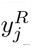 which is also the mean of 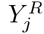 and its standard deviation 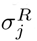. Schluter et al. (1997) showed that 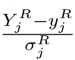 follows a *t*-distribution with *r* + 1 degrees o freedom. Namely, its density is:

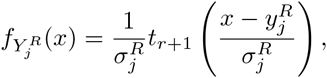

where *t*_*r*+1_ denotes the density of a *t*-distribution with *r* + 1 degrees of freedom:

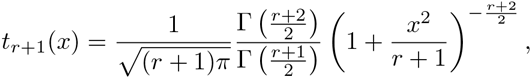

and Γ is the gamma function.
- As it is implemented in the ape R-package (Paradis et al. 2004), the Restricted Maximum Likelihood method (*REML*) reconstructs the ancestral states in a very similar way as *ML*. It first estimates the variance of the **BM** model. Next, the reconstructed ancestral states are those maximizing the likelihood under the **BM** model with the estimated variance. The relationship between *ML* and *REML* reconstructions is discussed in more details below. As for *ML*, 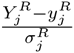 follows a *t*-distribution with *r* + 1 degrees of freedom.
- *PIC* is based on the Felsenstein’s Phylogenetic Independent Contrasts method (Felsenstein 1985). It recursively reconstructs the states of the ancestral nodes by averaging those of their children with weights depending on branch lengths. With *PIC*, the reconstructed state of a node only depends on those of its descendants. The confidence intervals are computed by using the expected variances under the model. They only rely on the tree (not on the leaf states). For any internal node *j, PIC* provides a reconstructed state 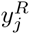, which also stands for the mean of 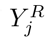, and the standard deviation 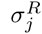 of 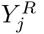. The random variable 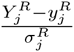 follows a standard Gaussian distribution.
- *GLS* stands for Generalized Least Squares method (Martins and Hansen 1997; Cunningham et al. 1998; Martins 1999). *GLS* reconstructs an ancestral state as a linear combination of those of the extant leaves according to the state variance-covariance structure under a given evolution model. By construction, *GLS* provides the best linear unbiased prediction of the ancestral states of a character evolving under this model (Martins and Hansen 1997). We consider only *GLS* based on the variance-covariance arising from a **BM** model. Here again, the confidence intervals are computed by using the expected variances under the model, thus only depend on the tree. For any internal node *j*, the random quantity 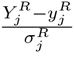 follows a standard Gaussian distribution and 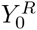 has the degenerate distribution at the reconstructed state of the root.

#### 2.1.2 Parsimony-based methods

There are two main kinds of parsimonious approaches dealing with quantitative characters: linear parsimony (Swofford and Maddison 1987; Maddison and Maddison 1992) and squared parsimony (*SP*) (Maddison 1991). The first one reconstructs the unknown states of the character by minimizing the sum of the absolute differences between the state of a node and that of its direct ancestor. The second one proceeds in the same way but it considers squared differences in place of absolute ones. Since, according to Butler and Losos (1997), linear parsimony often results in many equally parsimonious reconstructions and squared parsimony gives more relevant results, we keep only *SP* for our study. Unlike the methods of Section 2.1.1, *SP* does not provide any probability distribution for the reconstructed states.

### 2.2 Relations between methods

All these reconstruction methods are strongly related to the Maximum Likelihood reconstruction, thus one with another. These relations were already stated here and there, sometimes without justification. We recall them and give references or elements of proofs.

#### 2.2.1 SP

Minimizing the squared parsimony cost is equivalent to maximizing the log-likelihood under a **BM** model with a flat initial distribution for the root state and with all the branch lengths set to any constant value, see (Schluter et al. 1997; Maddison 1991) or Equation (1). It follows that any function computing *ML* may be used to compute *SP*. One just needs to make all the branch lengths equal before calling it.

#### 2.2.2 REML

Methods *ML* and *REML* only differ in the fact that the variance of the **BM** model and the ancestor states are simultaneously estimated by maximum likelihood with *ML*, while *REML* first estimates the variance of the **BM** model and next the ancestral states. If the root state follows an improper flat distribution then the term “log (*f*_*Z*0_(*z*_0_))” vanishes from Equation (1). Finding the ancestral states maximizing the log-likelihood just relies on finding the unknown values of the vector *z* minimizing 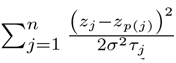. This does not depend on *σ*. Consequently, *ML* and *REML* do provide the same reconstructed states.

#### 2.2.3 GLS

Martins and Hansen (1997) state that *GLS* with the Brownian variance-covariance structure reconstructs the same ancestral states as *ML*. We provide a detailed proof of this fact in Appendix A.

#### 2.2.4 PIC

Maddison (1991) proved that *PIC* and *ML* reconstruct the same state for the root. Note that this only holds for the root.

#### 2.2.5 Totally equivalent?

*ML, REML* and *GLS* all reconstruct the same ancestral states. Nevertheless, they still have to be distinguished since they differ in terms of reconstructed distributions. Both *ML* and *REML* return *t*- distributions with *r* + 1 degrees of freedom and the same mean but with different variances, while *GLS* provides a Gaussian distribution with the same mean as *ML* and *REML* but with another variance.

### 2.3 Implementation

Since former implementations of *ML*, *REML* and *GLS* were based on numerical optimization algorithms, they did not always converge to global optimums. We used Equations (A5) of Appendix A to reconstruct ancestral states following these methods with exact matrix computations. The resulting R-function reconstruct is part of the **ape** R-package since version 3.2.

## 3 Assessing the performances of the reconstruction methods

We shall study the performances of the methods when the character is under directional evolution. To this aim, we assume that it evolves following an **ABM** model with variance *σ*^2^, drift *µ* and with the degenerate probability density at a given value *z*_0_ for the root state. We start this section by giving the conditional law of an internal state given those of the leaves under such a model. Next we propose an evaluation protocol using this conditional law. The Energy distance is strongly related to this protocol and allows us to compare between distributions and/or single values in a consistent way. Finally, we show that the conditional expectation of an ancestral state given the leaves is, in a sense, the best reconstruction possible and we study its relation with the state inferred by *ML*/ *REML*/ *GLS*.

### 3.1 Ancestral distributions conditioned on the leaf states

Let us assume here that the character evolution follows an **ABM** model (*z*_0_, *σ*^2^, *µ*). We put *Z_i_* for the random variable of the state *i* and *Z*, *Z*^(*a*)^ and *Z*^(*l*)^ for the random vectors *^t^*(*Z*_1_, …, *Z_n_*), *^t^*(*Z*_1_, …, *Z_r_*) and *^t^*(*Z*_*r*+_1,…, *Z_n_*), corresponding to all the nodes except the root, the internal nodes excluding root, and the leaves, respectively. A set of node states *z*_0_,…,*z_n_* is organized as vectors *z*, *z*^(*a*)^ and *z*^(*l*)^ accordingly.

Let *U* = *^t^*(*U*_1_, *U*_2_, …, *U_n_*) be the random vector of increments (i.e. *U_i_* = *Z_i_* — *Z*_*p*(*i*)_). Under the **ABM** model (*z_0_,σ^2^,µ*), the vector *U* is Gaussian with density:

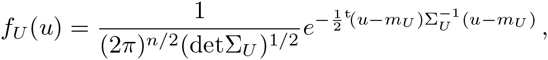

where *m_U_* is the expectation vector of *U* and Σ*_U_* its variance-covariance matrix which is diagonal since the coordinates of *U* are independent. We have 𝔼(*U_i_*) = *µτ_i_* and var(*U_i_*) = *σ*^2^τ*_i_* for all *i* ∈ { 1,…, *n* }.

In order to compute the joint law of the nodes, i.e. the law of the random vector *Z* = *^t^*(*Z*_1_, …, *Z_n_*), we remark that the vector *Z* is obtained from a linear transformation of *U*:

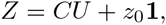

where *C* is a non-singular matrix. Since all its coordinates are affine combinations of the increments *U_j_*, which are independent Gaussian variables, the vector *Z* is still a Gaussian vector with density:

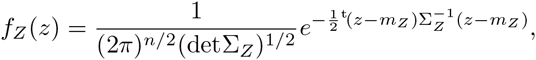

where *m_Z_* is the expectation vector of *Z* and Σ*_Z_* its variance-covariance matrix, namely

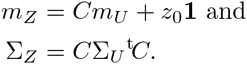

The matrix *C* is lower triangular (thanks to the nodes numbering) with diagonal entries all equal to 1 and other entries either equal to 0 or 1:

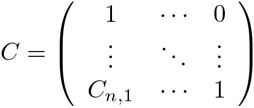

where 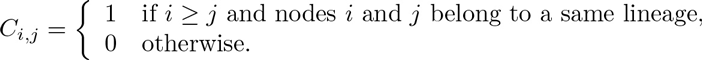

Putting *T_i_* for the time from root to node *i*, we have

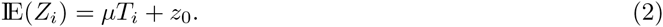

Since **ABM** models lead to Gaussian processes, the state distributions of all the nodes are multivariate normal. Let us compute the conditional joint law of the internal nodes given the leaf states *z*^(*l*)^, namely the law of *Y* = *^t^*(*Y*_1_, …, *Y_r_*) = (*Z*^(*a*)^|*Z*^(*l*)^ = *z*^(*l*)^). Let *m_a_* be the expectation vector of *Z*^(*a*)^ and ml that of *Z*^(*l*)^. Since the vector *Z* is a linear combination of the independent Gaussian increments *U_i_*, then any density *f_Z_*,*f*_*Z*^(*l*)^_ or *f*_(*Z*^(*a*)^_|*Z*^(*l*)^=*z*^(*l*)^) is multivariate Gaussian.

The variance-covariance matrix Σ*Z* of *Z* can be split according to *Z*^(*a*)^ and *Z*^(*l*)^:

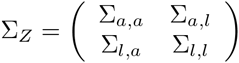

where Σ*_a,a_* is the variance-covariance matrix of *Z*^(*a*)^, Σ*_a,l_* is the covariance matrix between *Z*^(*a*)^ and *Z*^(*l*)^ and so on. The matrix Σ*_Z_* has the form *σ*^2^*K* where entry *K_ij_* is the time between the root and the most recent common ancestor of nodes *i* and *j* (Felsenstein 1973). We put 1 (resp. 1_*a*_ and 1_*l*_) for the n-dimensional (resp. r-
and (n — r)-dimensional) vector with all coordinates equal to 1.

With the decomposition of matrix Σ*_Z_*, the random vector *Y* is Gaussian with density *N* (*m_Y_, Σ_Y_*), where

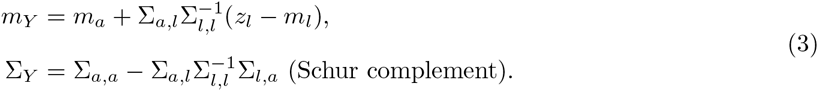

#### Remark 1.

*There exists a matrix M only depending on the tree such that Σ_Y_ = σ^2^M.*

*Proof*. From its definition, the remark holds for the matrix Σ*_U_*. It straightforwardly follows that it holds for Σ*_Z_*, thus for both Σ*_a,a_*, Σ*_a,l_*, Σ*_l,l_*, Σ*_l,a_* and finally for Σ*_Y_*.

In order to compute the conditional law of an ancestral state *i* given the leaves, we have to sum over all the other internal states. Since the marginals of a Gaussian vector are still Gaussian, we eventually get that *Y_i_* = (*Z_i_*|*Z*_*r*+1_ = *z*_*r*+1_, …, *Z_n_* = *z_n_*) follows an univariate Gaussian distribution 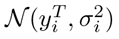, where 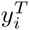 is the *i*^th^ coordinate of vector *m_Y_* and 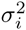 is the *i*^th^ diagonal entry of the variance-covariance matrix Σ*_Y_*.

Below, we use the ancestral distribution of the ancestral state *i* given the leaves as reference when comparing with its reconstructions. This conditional distribution is referred to as the *theoretical distribution* of state *i*. We put 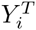 for the corresponding random variable.

### 3.2 Evaluation protocol

#### 3.2.1 For reconstructed states Absolute bias

In a simulation context, the relevance of a reconstructed state is generally assessed by measuring its distance from the corresponding simulated ancestral state (Butler and Losos 1997; Martins 1999; Oakley and Cunningham 2000; Webster and Purvis 2002). This distance accounts for the reconstruction error. Remark that, in order to make sense, the distances between the reconstructed and simulated states have to be averaged over a large number of simulations.

The theoretical distributions derived in the previous section may be used to improve the assessment of reconstructed states. Let us consider an evolution *z* simulated under an **ABM** model. A reconstruction method only takes into account the leaf states *z*_*r*+1_, …, *z_n_*. On the other hand, Section 3.1 gives us the conditional distribution of an ancestral state *i* given *z*_*r*+1_, …, *z_n_* under the simulation model. Intuitively, this distribution would be asymptotically observed by running an infinite number of simulations and by keeping only those with leaf states *z_r+_1, …, z_n_*. The distance expectation between 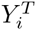 and the reconstructed state is exactly the conditional expectation of the reconstruction error on state *i* given the leaf states.

This suggests to replace the standard evaluation procedure by the following protocol. Being given a distance d,

1. simulate an evolution *z* under an ABM model;
2. retain only the leaf states *z*_*r*+1_, …, *z_n_*;
3. for all nodes *i*, compute from *z*_*r*+1_, …, *z_n_*:

- the reconstructed state 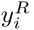
- the conditional distribution of state *i* given *z*_*r*+1_, …, *z_n_* under the simulation model (i.e. the distribution of 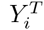);
4. for all nodes *i*, compute 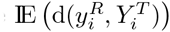

This protocol ensures that the leaf states are well sampled from their probability distribution under the simulation model. It follows that averaging 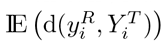, which is conditioned on the leaf states, over all the simulations do converge to the expected reconstruction error on state *i* under the simulation model.

In the standard evaluation scheme, the distance d between a reconstructed state 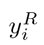 and the corresponding simulated state 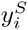 is generally measured in terms of bias 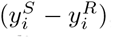, absolute bias 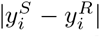 or squared bias 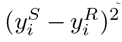. Let us compute the expectations of these distances between a reconstructed state and the random variable 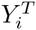 following the theoretical distribution 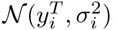. They are

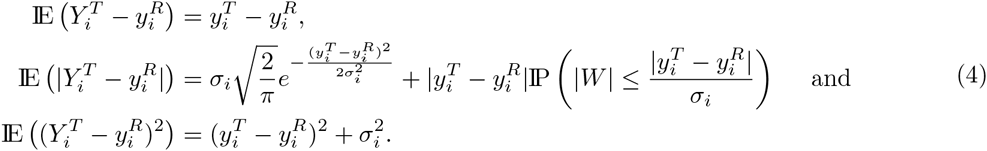

where *W* stands for the standard Gaussian variable. Although all these measures are suitable to compare the random variable 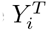 with the reconstructed state 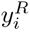 they do not take into account the same amount of information from the distribution of 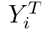. The bias is only based on its mean, the squared bias uses its mean and variance while the absolute bias takes into account both its mean, variance and the normality of the distribution. This point somehow supports the choice of this last distance.

#### 3.2.2 For reconstructed distributions Energy distance

Assessing the relevance of the uncertainty distributions provided by the reconstruction methods could also be done by considering the simulated ancestral states. But one expects more efficiency by considering the theoretical distributions. Adapting the above protocol to this case could be done by considering, for all nodes *i*, the expectation 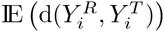, where 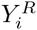 follows the distribution provided by the method for *i*, which is also conditioned on the leaf states. A major drawback here is that the above expectation is not a good measure of the similarity between two probability distributions. In particular, it is not equal to zero when 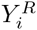 and 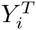 are identically distributed (and not degenerate).

On the other hand, there exist various distances for comparing two probability distributions. Among them, the so-called Energy distance is strongly related to the evaluation protocol when d is the absolute bias. We will see that it offers us a consistent framework to compare states versus states, states versus distributions and distributions versus distributions.

Let *A* and *B* be two random variables and *F_A_* and *F_B_* their respective cumulative distribution functions. For convenience reasons, we write the distance between two distributions as the distance between two random variables following them. There are two equivalent ways to define the *Energy distance* (E-distance) between *A* and *B* (Sz´ekely and Rizzo 2013):

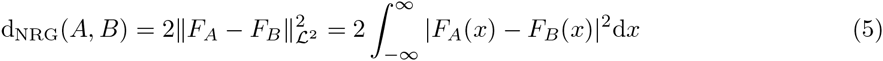

and

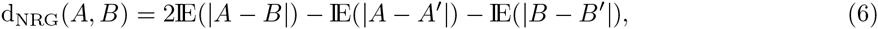

where *A′* and *B′* are independent and identically distributed copies of *A* and *B* respectively.

A distance between distributions can be used for comparing a single value against a distribution (or even two single values), just by considering the degenerate distribution(s) at the single value(s).

Let us start by checking the behavior of the E-distance when comparing two single values. Assuming that *A* and *B* follow degenerate distributions at *a* and *b* respectively, we have that

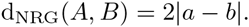

In plain English, the E-distance between two degenerate distributions is twice the absolute bias between the corresponding values.

Now if *A* follows the degenerate distribution at *a* and *B* follows a Gaussian distribution with variance 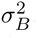, we have to compute the expected absolute value of Gaussian variables, whose formula is recalled in Appendix B, Equation (B1). Thus the E-distance becomes:

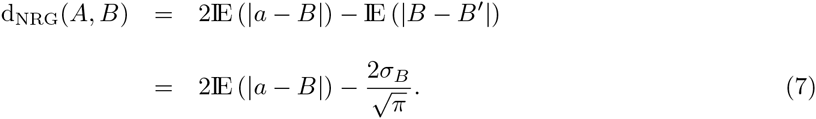

The E-distance between a reconstructed state and the corresponding theoretical distribution is twice the expectation of the absolute bias between the state and a random variable following the theoretical distribution, minus a correcting term. Up to this term, averaging the E-distances between the reconstructed states and the theoretical distributions is the same as applying the evolution protocol with the absolute bias.

Finally, Equation (6) shows that the Energy distance between two random variables following general distributions is twice the expectation of the absolute bias between them, minus two terms which somehow accounts for their respective dispersions.

In conclusion, the E-distance is strongly related to the protocol of Section 3.2.1 when evaluating reconstructed states or reconstructed distributions with the absolute bias.

The protocol eventually used in our comparisons is that of Section 3.2.1 with the 4^th^ step replaced by 4. for all nodes *i*, compute 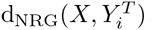.
where *X* is either the degenerate random variable of the reconstructed state at i or a random variable following the uncertainty distribution provided by the method under evaluation.

We show how to compute the E-distance of pairs of distributions involved in a theoretical vs reconstructed distributions comparison, i.e. Gaussian versus degenerate, Gaussian versus Gaussian and Gaussian versus Student, in Appendix B (R-scripts available on request).

The more usual Kolmogorov-Smirnov distance is actually harder to interpret when degenerate distributions are involved. In particular the Kolmogorov-Smirnov distance between two degenerate distributions is always 1 except if they are equal (Suppementary material). Supplementary Figures S3 and S4 display the Kolmogorov-Smirnov distances. Their general behavior is the same as with the Energy distance (Figures S1 and S2).

### 3.3 Optimal reconstruction

Let us assume that the character follows an **ABM** model (*z*_0_,*σ*^2^,*µ*). Being given a set of leaf states and under the model, reconstructing the state of *i* with the mean 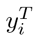 of its theoretical distribution leads to the smallest expectation error in terms of any standard distance (Equation 4) and of E-distance (Equation 7). The argument is similar as that of (Steel and Sz´ekely 1999). Namely, reconstructing with 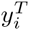 leads to expectations of bias, absolute bias and squared bias equal to 0, 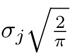 and 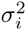 respectively, and to E-distance 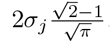. The mean 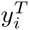 will be referred to as the *optimal reconstruction* of the state *i*. Remark that computing 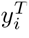 requires to know the parameters of the model of evolution. The optimal reconstruction can be determined in a simulation context but unfortunately not in a practical situation.

In the particular case of a **BM** model, the optimal reconstruction is strongly related to the state reconstructed by *ML/REML/ GLS*. Indeed, let us consider a **BM** starting at the grand mean 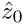, given by the first formula of Equation (A5) in Appendix A. By considering Equation (2) with *µ* = 0, the partial mean vectors *m_a_* and *m_l_* are equal to 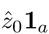 and 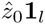 respectively. It follows that the second equation of (A5), which gives the ancestral reconstructed states, and the first equation of (3), which gives the conditional means, are identical. In short, if *ML/REML/ GLS* infers the **“**real**”** state of the root, it reconstructs the whole tree in an optimal way.

## 4 Results and discussion

### 4.1 Simulation protocol

Although the simulation model of evolution has three parameters (*z*_0_, *σ*^2^, *µ*), we only vary the last two ones. We keep the root state *z*_0_ fixed since it just translates the whole process and does not influence the methods performance. In order to assess their accuracy we simulate the evolution of a quantitative character along the Pleistocene planktic Foraminifera phylogenetic tree (Webster and Purvis 2002), given in Figure 1, which starts from *z*_0_ = 100 at the root and evolves under ABM models with various drifts and variances (21 values for parameter µ, ranging from —10 to 10 and 15 values for parameter *σ*, ranging from 0.01 to 20).

**Figure 1:**
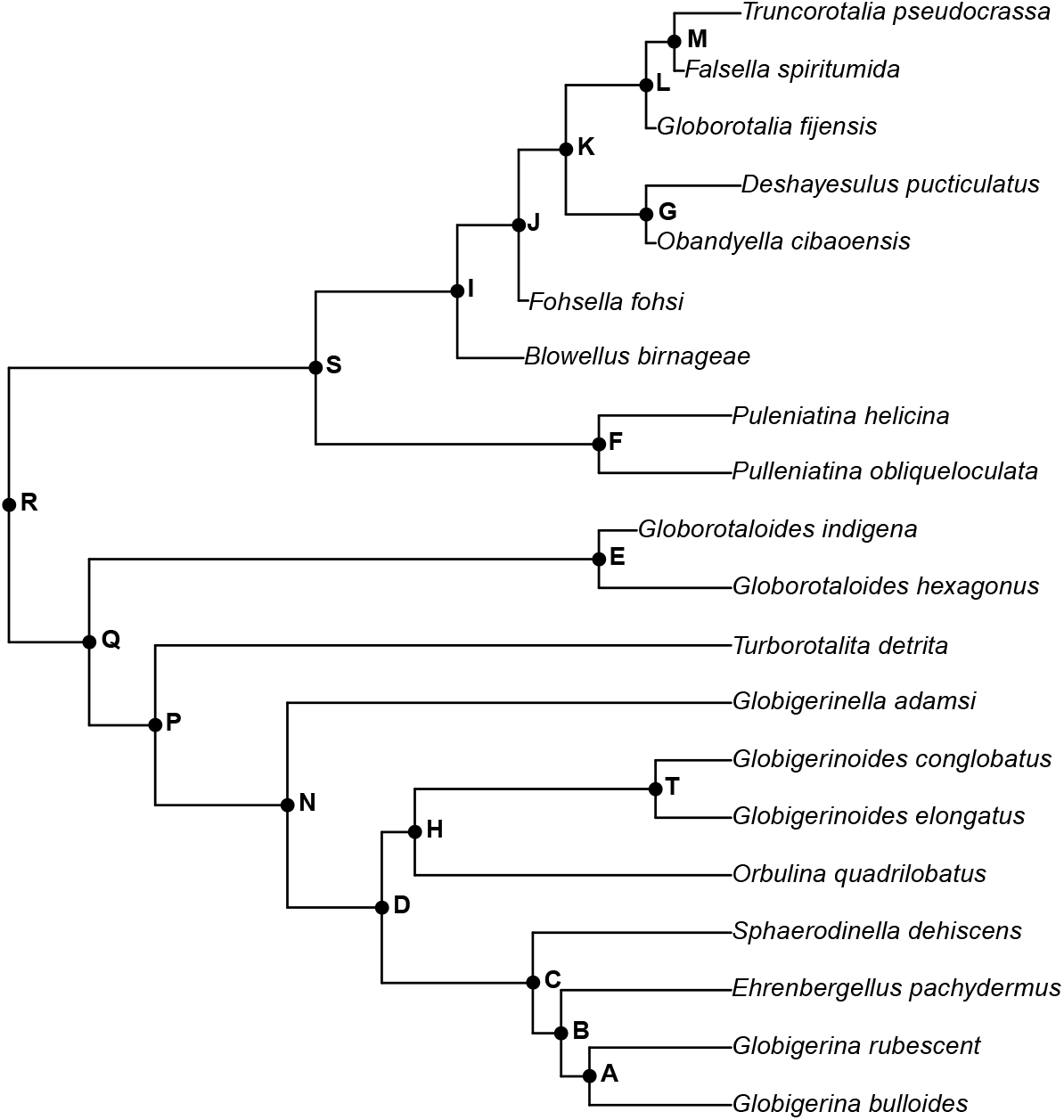
Pleistocene planktic Foraminifera phylogeny (Webster and Purvis 2002) on which the simulations runs.

For each parameter set, we run 500 simulations from which we retain only the leaf states. We apply the evaluation protocol of Section 3.2 on the reconstructed states and on the reconstructed distributions provided by *PIC* from the function ace, and by *ML, REML, GLS* and *SP* from the function reconstruct of the ape R-package. The theoretical distribution of each ancestral state under the simulation model is then compared first, with the reconstructed states and second, with the corresponding reconstructed distributions, in both cases in terms of E-distance. These distances are finally averaged over all the simulations in order to compare the performances of the methods.

### 4.2 Single reconstructed states

We first evaluate the methods accuracy with regard to the ancestral states they provide. Since *ML*, *REML* and *GLS* return the same inferred states, they have the same accuracy which is compared with that of *PIC*, *SP* and with the optimal one.

#### 4.2.1 Lower bounds of reconstruction errors

Since - except in degenerate cases - there is no unique ancestral states configuration leading to the leaf states from which we infer, reconstructing an ancestral state with a single value always comes with a certain probability of error. A counterpart of this fact is that the E-distance between a reconstructed state and its theoretical distribution is positive. From Section 3.3, the smallest E-distance which can be obtained with a reconstructed state 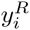 comes by setting 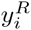 equal to the mean of the theoretical distribution 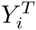. It leads to a E-distance 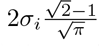 with 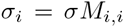 where *M_i,i_* only depends on the tree (Remark 1), whatever the drift µ and the root state *z*_0_. To sum up, being given the parameters (*z*_0_, *σ*^2^, *µ*) of the model, the optimal reconstruction provides a lower bound for the E-distances which depends linearly on the variance of the model, but neither on its trend nor on its initial value. It is represented by red lines in Figures 2 and 3.

**Figure 2:**
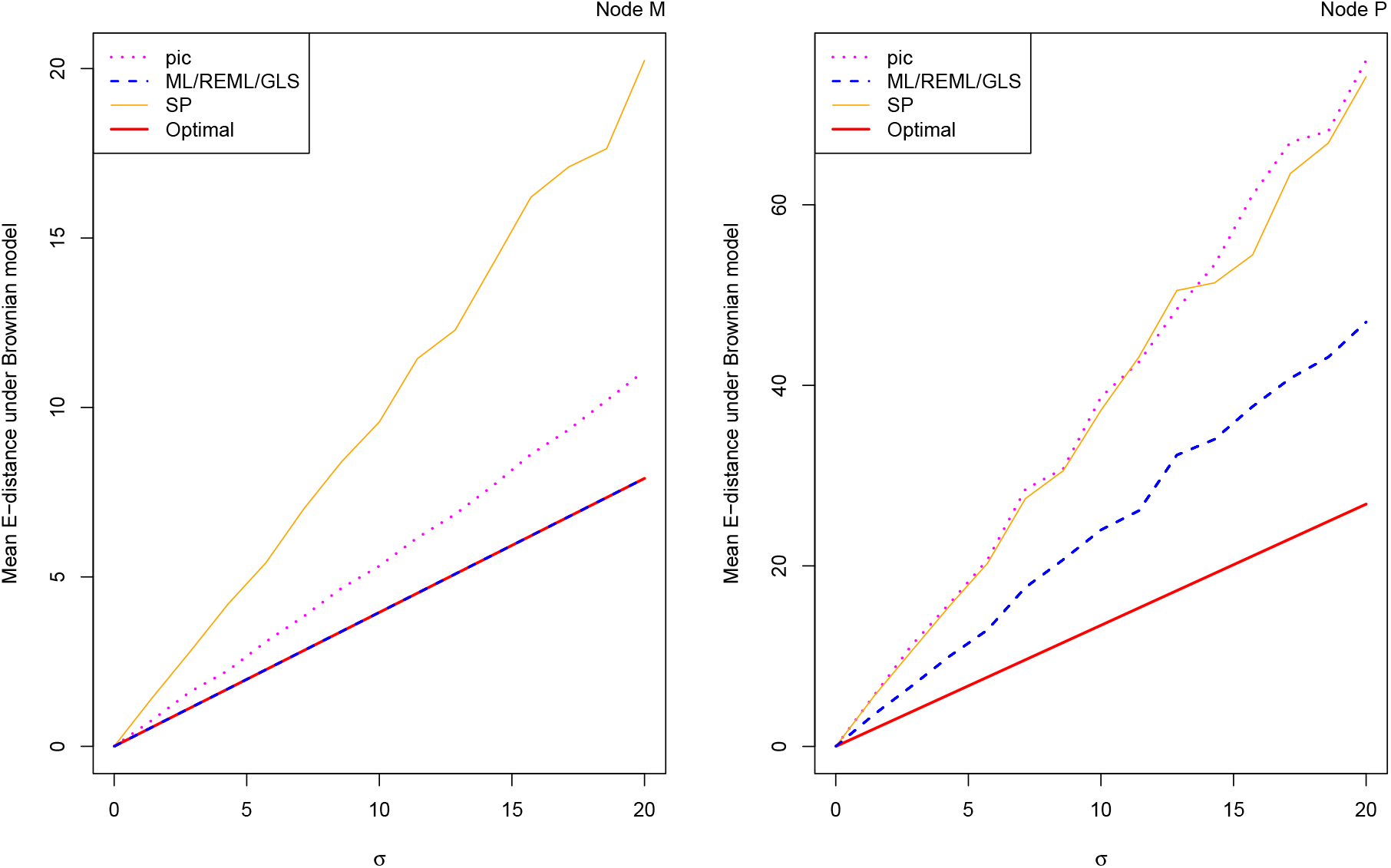
Mean Energy distance between the state reconstructed by *PIC*, *ML*/*REML*/*GLS* and *SP* and the corresponding theoretical distribution from **BM** models versus the parameter σ for the nodes M and P.

**Figure 3:**
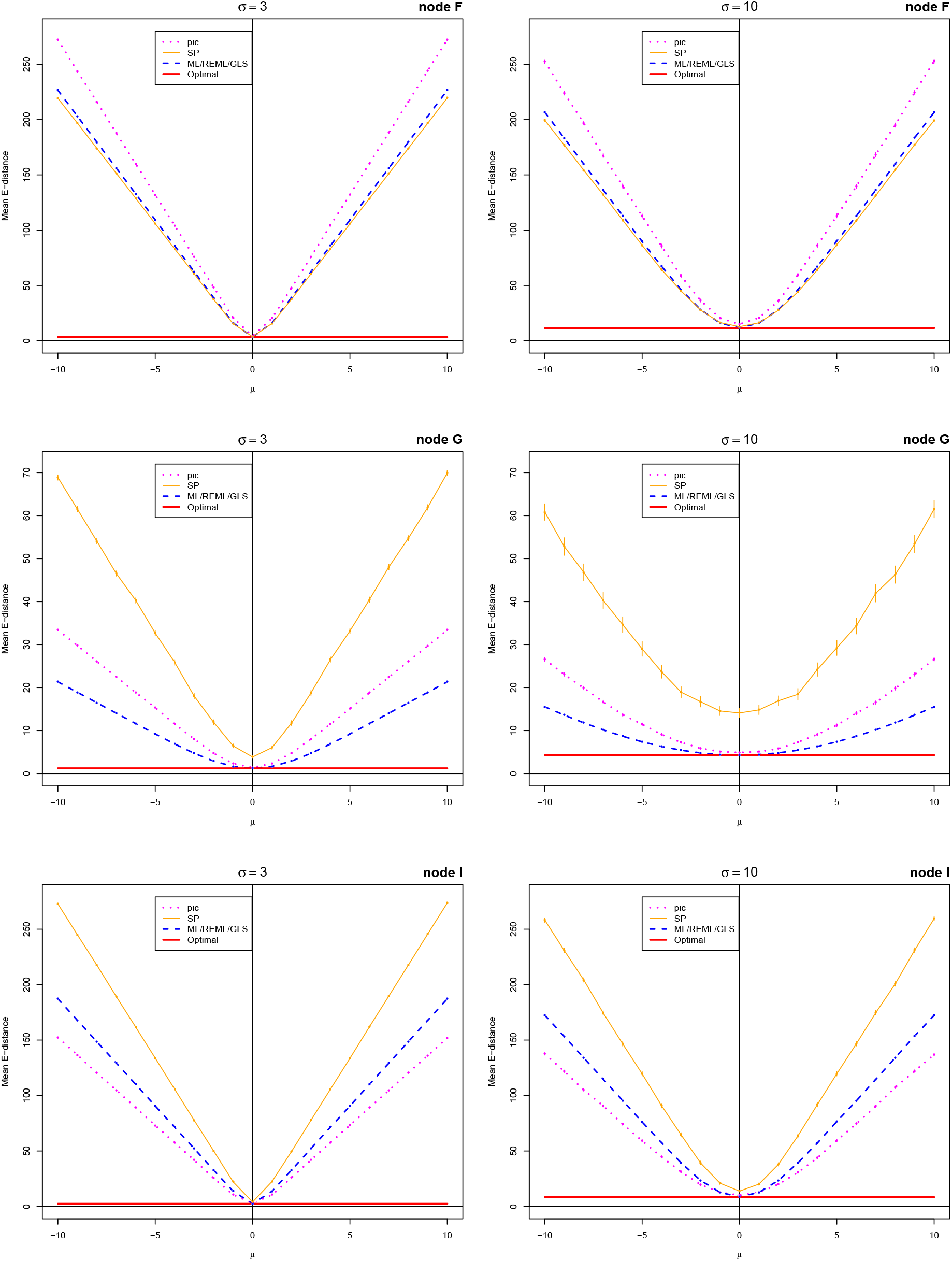
Mean Energy distance between the state reconstructed by *PIC*, *ML*/*REML*/*GLS* and *SP* and the corresponding theoretical distribution from **ABM** models with σ equal to 3 and 10 versus the parameter µ for the nodes F, G and I.

Asking whether a method achieves the optimal reconstruction, at least when the character follows a **BM** model, is a natural question. *ML/REML/ GLS* sounds like a good candidate for that, since it reconstructs the optimal state for any node as soon as the inferred root state matches *z*_0_ under a **BM** model (Section 3.3). We do observe that the states inferred by *ML/REML/ GLS* are indistinguishable from the optimal ones for nodes A, B, G, J, K, L, M and T but not for the other nodes. These two situations are shown in Figure 2, which displays the results of nodes M and P. The fact that *ML/ REML/ GLS* are not optimal for some nodes always comes from an inaccurate estimation of the root state. Remark that despite this inaccurate root estimation, *ML/REML/GLS* may still be almost optimal for some of the nodes.

#### 4.2.2 Influence of the simulation parameters

The smaller the parameter *σ* of a **BM** model, the more accurate the reconstructions of all the methods (Figure 2). Basically, as *σ* decreases, all the states of the tree (both ancestral and tips) get closer to one another, which makes the reconstruction easier.

Another general observation is that, under an **ABM** model, all the methods perform better as the drift µ is close to 0 (Figure 4). This was expected since this situation is close to a **BM** model which is the assumption underlying all the methods but *SP*. The influence of µ is very strong for nodes A, B, C, D, F, H, I, N, P, Q, R, S and T (E-distances from different µ are far from each other) but is much less striking for nodes E, G, J, K, L and M for which plots sometimes overlap. Figure 4 displays the E-distances of the nodes **D** and **J.**

**Figure 4:**
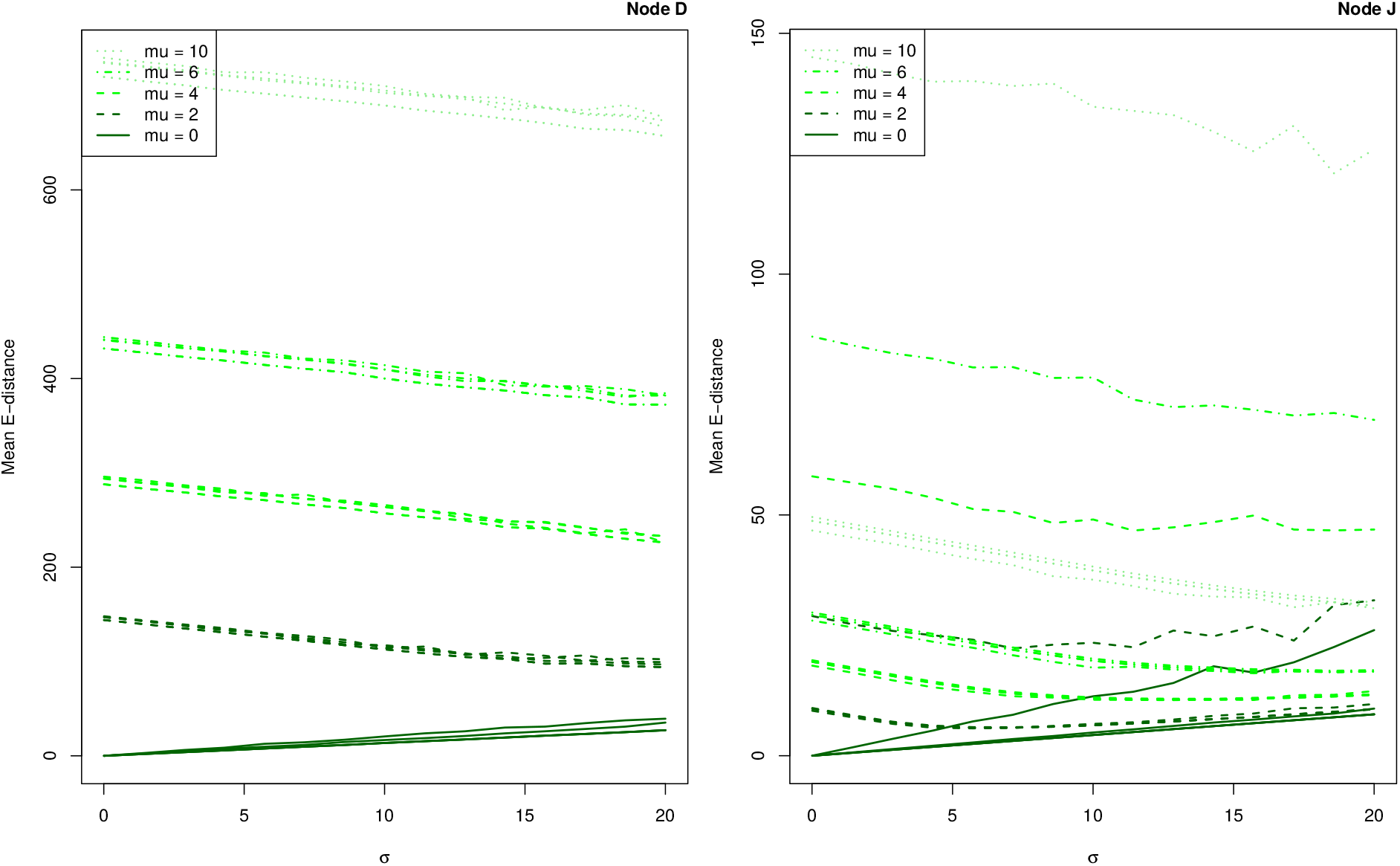
Mean Energy distance between the state reconstructed by *PIC*, *ML*/*REML*/*GLS* and *SP* and the corresponding theoretical distribution from **ABM** models with different values of µ versus the parameter σ for the nodes D and J.

#### 4.2.3 Methods comparison

The methods performances are very close to one another for some of the nodes. This is basically the case for the root for which *ML/REML/ GLS* and *PIC* infer the same reconstructed state, but this is also observed for nodes A, B, C, D, H, N, Q and T. For this reason, we add error bars representing 95%-confidence intervals for the mean E-distance in Figure 3. Whenever the error bars do not overlap, the method corresponding to the lower curve has a significantly better performance than that of the upper one, according to the Student’s t-test for paired samples with *α* = 5%.

Under a **BM** model, corresponding to data simulated with *µ* = 0, *ML/REML/ GLS* provides the most accurate reconstruction for the nodes displayed in Figures 2 and 3. This is actually observed for all the nodes of the tree. Supplementary Figure S1 displays the plots of Figure 3 for all the nodes. Still under a **BM** model, the relative ranking of *SP* and *PIC* depends on the node. Indeed, for nodes P and Q, performance ranking of *SP* and *PIC* varies with *σ*, while for nodes A, B, D, F, H, K and S, *SP* is more accurate than *PIC*, the opposite being observed for the remaining nodes. Figure 2 displays an example where *PIC* is better than *SP* and one where the most accurate method changes with *σ*.

Comparing the methods from data simulated with a significant drift is more puzzling. Though *ML/REML/GLS* leads to the smallest E-distance in general, this behavior is not observed with nodes E, I, J and S for which the minimum E-distance is achieved by *PIC* and with node F for which it is achieved by *SP*.

### 4.3 Reconstructed distributions

Let us evaluate the relevance of the distributions provided by the methods for the reconstruction uncertainty. The methods are now assessed with the E-distance between the theoretical distribution and the reconstructed one which is either Gaussian (*PIC* and *GLS*), Student (*ML* and *REML*), or degenerate (*SP* and *GLS* at the root). As some of the methods could possibly provide a reconstructed distribution matching exactly the theoretical one, we no longer have a lower positive bound of these E-distances (the counterparts of the **“**optimal**”** red lines of Figure 3 are the abscissa axis in Figure 5). Since they are Student instead of Gaussian, the distributions provided by *ML* and *REML* can not perfectly match the theoretical ones. However, though *GLS* and *PIC* could conceivably lead to a perfect match with the theoretical distribution, their performances are always significantly – we keep plotting errorbars – overcome by *ML* and/or *REML*.

**Figure 5:**
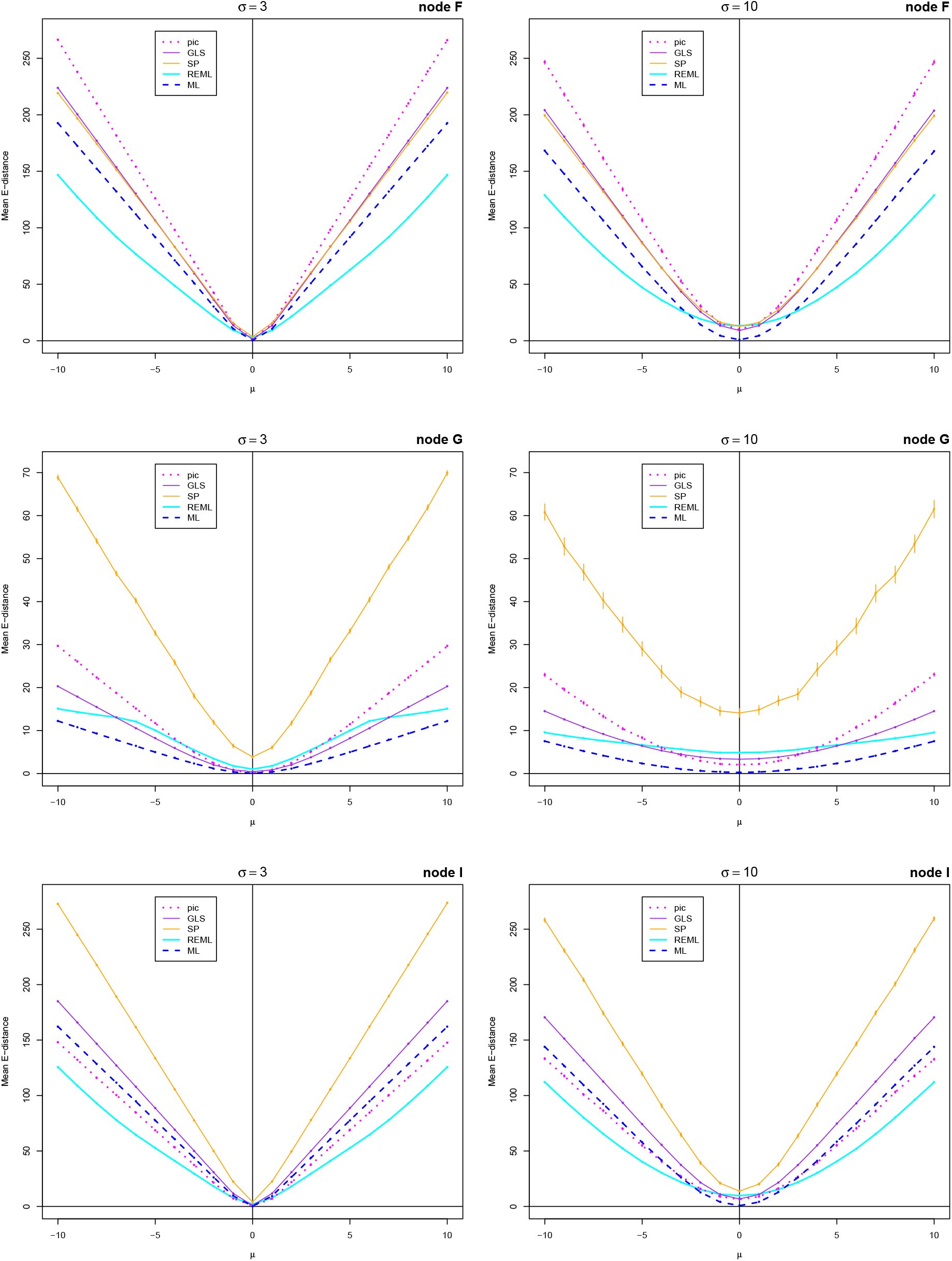
Mean Energy distance between the reconstructed distribution from *PIC*, *ML*, *REML*, *GLS* and *SP* and the corresponding theoretical distribution from **ABM** models with σ equal to 3 and 10 versus the parameter µ for the nodes F, G and I.

Considering reconstructed distributions rather than single values is expected to change the E-distances. The most notable difference is that *ML*, *REML* and *GLS* are no longer equivalent and can now be compared one another as well as with *PIC* and *SP*. Note that the E-distances are the very same in the case of *SP*, since this method only provides a reconstructed state. We observe an improvement on the *PIC* plots for node G between Figures 3 and 5. The same can be observed for node M, but the curves are nearly indistinguishable for all the other nodes. Like in the case of *PIC*, there is no observable change for *GLS*, except for nodes G and M, where its performance is slightly improved. On the contrary, the distributions provided by *ML* and *REML* do lead to lower E-distances than those obtained from the single reconstructed states. A general observation is that *ML* is significantly better than all the methods when the drift µ is close to 0 (i.e. when the model is close to **BM**). As the drift increases, *ML* remains the best method for nodes G, L and M but it tends to be overcome by *REML* for all the other nodes. Supplementary Figure S2 shows the plots of Figure 5 for all the nodes of the tree.

### 4.4 Discussion

As expected, we observe that all the methods perform better when the character evolution follows a **BM** model. Under an **ABM** model, the reconstruction accuracy decreases linearly with the drift intensity. Our simulations show that the reconstruction methods may lead to spurious results if the character does not evolve under neutrality, like it was observed with fossil data in (Webster and Purvis 2002).

By construction, the theoretical distributions obtained from the simulation model reflect the real uncertainty of the character reconstruction. Thus one expects from the distributions provided by a reconstruction method to approach the theoretical ones or, at least, to be more informative that single reconstructed states. Unfortunately, the reconstructed distributions provided by PIC and GLS are generally farther from the theoretical ones than the degenerate laws at their reconstructed states. The way in which they are computed is quite general in the sense that it does not depend on the leaf states. On the contrary, the distributions provided by *ML* and *REML* are actually closer to the theoretical ones than the degenerate laws at the reconstructed states. This indicates that these distribution may be relevant with regard to the inherent reconstruction uncertainty.

To summarize the results of our comparison, we observe that *ML* is the most accurate – or at least among the most accurate – each time the character evolution is close to neutral. As the drift intensity increases and although *ML* and *REML* still often provide the most accurate reconstructions, they are sometimes overcome by PIC or even SP. Overall, the methods do not deal well with drift. Their comparison is inconclusive for characters under directional evolution. This calls for the development of ancestral state reconstruction methods able to take into account a trend on the character evolution.

## Acknowledgements

We thank Bastien Bousseau, Pierre Pontarotti and Laurence Reboul for their careful readings and their helpful remarks and suggestions.

## Appendix

## A Equivalence between *GLS* and *ML*

Let *Z_i_* be the random variable of the node state *i*. We put *Z*, *Z*^(*a*)^ and *Z*^(*l*)^ for the random vectors *^t^*(*Z*_1_, …, *Z_n_*), *^t^*(*Z*_1_, …, *Z_r_*) and *^t^*(*Z*_*r*+1_,…, *Z_n_*), corresponding to all the nodes except the root, the internal nodes excluding root, and the leaves, respectively. A set of node states *z*_0_, …, *z_n_* is organized as vectors *z*, *z*^(*a*)^ and *z*^(*l*)^ accordingly.

In order to explain why *GLS* and *ML* are equivalent, we shall consider two different expressions of the likelihood under the assumptions of *ML*. In particular, *ML* assumes that the character evolution follows a **BM** model with variance *σ^2^*. The probability density of a vector *z*_0_, …, *z_n_* can be written either

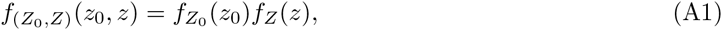

or

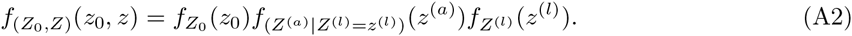

Since the vector *Z* can be expressed as a linear transformation of the independent Gaussian increments *Z_j_*-*Z*_*p*(*j*)_, both *f_Z_*, *f*_*Z*(*l*)_ and *f*(_*Z*_^(*a*)^|*Z*^(*l*)^=*z*^(*l*)^) are multivariate Gaussian densities. The variance-covariance matrix Σ*_Z_* of *Z* can be split according to *Z*^(*a*)^ and *Z*^(*l*)^:

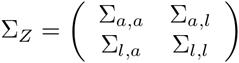

where Σ*_a,a_* is the variance-covariance matrix of *Z*^(*a*)^, Σ *_a,l_* is the covariance matrix between *Z*^(*a*)^ and *Z*^(*l*)^ and so on. The matrix Σ*_Z_* has the form *σ^2^K* where entry *K_i,j_* is the time between the root and the most recent common ancestor of nodes *i* and *j* (Felsenstein 1973). We put 1 (resp. 1_*a*_ and 1_*l*_) for the n-dimensional (resp. *r*-and (*n* - *r*)-dimensional) vector with all coordinates equal to 1. Since *Z* follows the multivariate normal distribution *N*(*z*_0_1, Σ_Z_), the marginal and conditional random vectors *Z*^(*l*)^ and (*Z*^(*a*)^|*Z*^(*l*)^ = *z*^(*l*)^) follow the multivariate normal distributions *N*(*z*_0_ 1_l_, Σ_l,l_) and 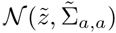 respectively, where

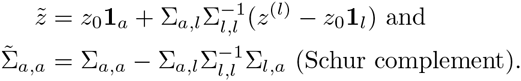

Under the *ML* assumptions, *f*_*Z*_0__ is the improper flat density, thus its logarithm just vanishes in the computation of *log*(*f*(*Z*_0_,*Z*)^(*z*_0_, *z*)^) with Equations (A1) and (A2). On the one hand, from Equation (A2), the vector of partial derivatives of *log*(*f*(_*Z*_0_,*Z*_)^(*z*_0_,*z*)^) with respect to the internal states *Z*^(*a*)^ is proportional to the vector

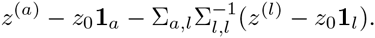

On the other hand, from Equation (A1), the partial derivative of *log*(*f*(*Z*_0_,*Z*)^(*z*_0_, *z*)^) with respect to the root state *z*_0_ is proportional to

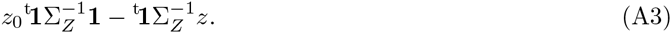

The maximum likelihood estimates 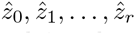 of internal states with respect to the vector leaf states *Z*^(*l*)^ may basically be obtained by solving the system of linear equations:

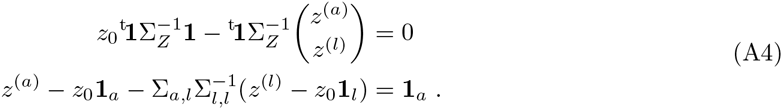

Let us get a simpler form for the first equation of (A4). The inversion formula for block matrices gives us that

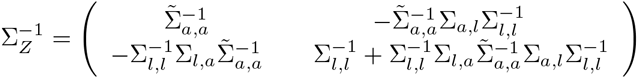

It follows that Expression (A3) can be rewritten as

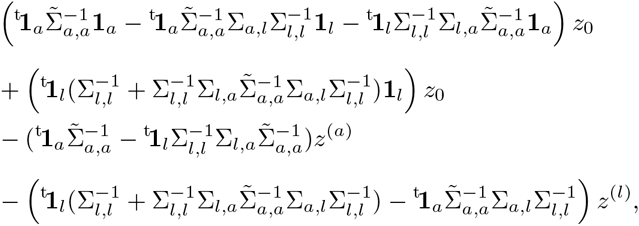

in which, substituting *Z*^(*a*)^ according to the second equation of (A4), leads to

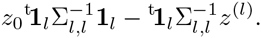

Finally, the estimates 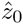 and 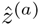 maximizing the log-likelihood with respect to the vector of leaf states *Z*^(*l*)^ satisfy:

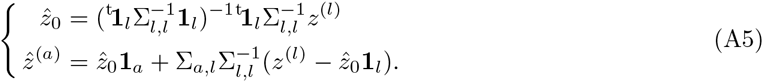

These formula are the same as those computing the *GLS* reconstruction (Martins and Hansen 1997; Cunningham et al. 1998; Martins 1999) in which zˆ_0_ is called the *grand mean*.

## B Energy Distance

Let *A* and *B* be two random variables and *F_A_* and *F_B_* their respective cumulative distributions.

- If both *A* and *B* follow degenerate distributions at *a* and *b* respectively, then

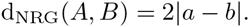
- If *A* follows a normal law 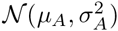 and *B* a degenerate distribution at b, then, since for a standard Gaussian random variable *W*

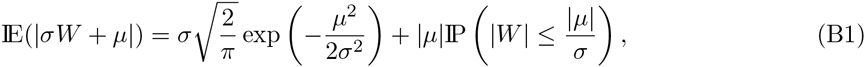

we have

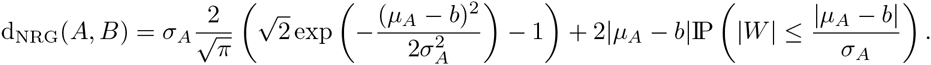 If *µ_A_* = *b*, the Energy distance is equal to 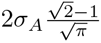 thus increases with *σ_A_*. For any fixed *σ_A_* = 0, the Energy distance goes to infinity as |*µ_A_* - *b|* becomes larger while for any fixed distance |*µ_A_ - b|* = 0, it goes from 2|*µ_A_* - *b|* to infinity as *σ_A_* goes from 0 to infinity.
- Let us assume that *A* and *B* follow a degenerate distribution at *a* and a Student law 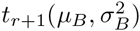 respectively. For the Student random variable *W* with (r + 1) degrees of freedom, we have

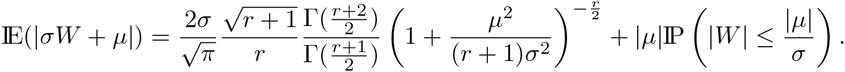 It is possible to compute directly IE(|*A - B|)* and IE(|A-*A’|)*, but not IE(|B - *B’|)* because the difference between two independent Student variables is not a Student variable. We thus rely on Expression (5) of the Energy distance:

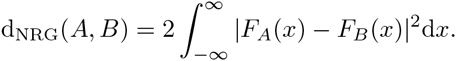

with 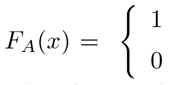 if *a ≤ x*, otherwise. and 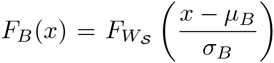, where *F_Ws_* is the Student cumulative distribution function with r + 1 degrees of freedom. The integral of (5) is approximated by 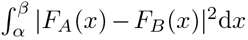, where 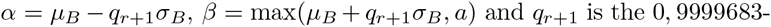 quantile for a Student variable with r + 1 degrees of freedom. The numerical computation of this last integral is performed by the Wynn’s Epsilon algorithm (Piessens et al. 1983).
- If both *A* and *B* are Gaussian variables, we get from Equation (B1) that

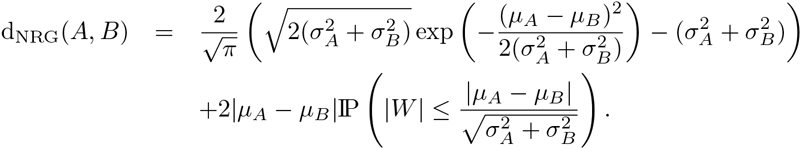
- Let us finally assume that *A* and *B* follow a normal distribution 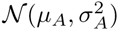 and a Student distribution 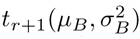 respectively. Hence again we have to rely on Expression (5). We approximate the integral with 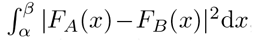, where 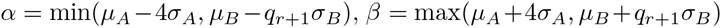 and q_r_+ | (resp. 4) is the 0, 9999683-quantile for a Student variable with r + 1 degrees of freedom (resp. for a standard Gaussian variable). The integral is computed via the Wynn’s Epsilon algorithm.

